# Exosome-derived lncRNA HOXA-AS3 promotes castration resistance and progression of prostate cancer via the miR-29b-3p/Mcl-1/STAT3 axis

**DOI:** 10.1101/2022.09.28.509879

**Authors:** Jie Teng, Yan Zhao, Limin Shang, Yang Li, Jian Zhang, Liang Zhu, Yegang Chen, Gang Li, Zhifei Liu, Mingfei Jia, Shaosan Kang, Haitao Niu, Yuanjie Niu, Qiliang Cai

**Affiliations:** Department of Urology, Tianjin Institute of Urology, the Second Hospital of Tianjin Medical University, Tianjin, 300211,China; Department of Endocrinology, the Second Hospital of Tianjin Medical University, Tianjin, 300211,China; Department of Urology, Beijing Friendship Hospital, Capital Medical University, Beijing, 100050, China; Department of Urology, Tangshan People’s Hospital, Tangshan, 063000, China; Department of Urology, North China University of Science and Technology Affiliated Hospital, Tangshan, 063000, China; Department of Urology, the Affiliated Hospital of Qingdao University, Qingdao, 266003, China

**Keywords:** Prostate cancer, androgen resistance, progression, exosomes, lncRNA HOXA-AS3

## Abstract

Prostate cancer is one of the leading causes of cancer-related deaths in men. While endocrine therapy is effective in the early stage of metastasis, significantly inhibits the progression of metastatic prostate cancer, most patients eventually develop CRPC. Tumor microenvironment are involved in the progression of prostate cancer as an “accomplice”, cancer cell–secreted exosomes were identified as crucial messengers can carry lncRNAs to participate in intercellular communication. we revealed PC-3-derived exosomes promote androgen resistance in LNCaP cells. HOXA-AS3 as a ceRNA of miRNA-29b-3p affects the proliferation and invasion ability of prostate cancer cells. A series of molecular experiments, cell experiments and clinical tissue verification experiments confirmed that HOXA-AS3 participates in the castration resistance and progression of prostate cancer through regulating the miR-29b-3p/Mcl-1/STAT3/Cytochrome C/caspases-9 pathway. Dysregulation of HOXA-AS3 is observed in many cancer types, and this study shows the importance of this lncRNA in controlling prostate cancer cell progression, thus highlighting it as a potential biomarker for inhibiting prostate cancer progression and a target for immunotherapy.

## Introduction

Prostate cancer (PCa) is the most common malignant tumor in the male reproductive system, with approximately 1.3 million new cases diagnosed worldwide every year (Sandhu S et al, 2021). While androgen-deprivation therapy (ADT) therapy is effective in the early stage of metastasis, significantly inhibits the progression of metastatic prostate cancer and greatly improves the 5-year survival rate and quality of life of patients, most patients eventually changed from ADPC to CRPC (Cornford P et al, 2017), which decreases survival rate and reduces quality of life of the patients significantly (Nelson WG et al, 2003). Therefore, it is crucial to explore the mechanism and development of hormone-independent prostate cancer.

Exosomes play an important role in cell-cell interactions in the tumor microenvironment (Zhang L et al, 2019), and involved in tumor progression, promoting the angiogenesis and migration of tumor cells during metastasis. Given the specific characters of exosomes, their use as a new platform for cancer therapy and as cancer biomarkers is promising. Many studies have shown that exosomes are involved in the development of prostate cancer. Han Q et al found that silencing SIRT6 by siRNA delivered through exosomes inhibited tumor growth and metastasis (Han Q et al, 2021). Overexpression of exosomal miR-1246 in PCa xenograft mouse model in vivo highlighted the tumor suppressive role of this miRNA suggesting its therapeutic relevance in managing aggressive PCa (Bhagirath D et al, 2018). Therefore, we speculate that exosomes play a significant role in the progression of androgen-resistant prostate cancer to androgen-dependent prostate cancer.

LncRNAs, are pervasive noncoding RNA classes with important biological roles that can regulate the expression of downstream target genes and participate in the progression and metastasis of multiple malignancies through different mechanisms, such as the recruitment of chromatin regulators and transcription factors (Prensner JR et al, 2013; Guo H et al, 2016; Wu XS et al, 2017; Liang Y et al, 2018). Previous individual lncRNAs in prostate cancer through in vitro cell line and in vivo xenograft models (Prensner JR et al, 2013; Guo H et al, 2016; Hua JT et al, 2018), SChLAP1, PCAT1, PCAT18, and PCAT19 were found to contribute to prostate cancer cell growth and aggressiveness. Growing evidence has shown the oncogenic role of long non-coding RNA HOXA-AS3 in the progression of several types of cancers, HOXA-AS3 facilitates the malignancy in colorectal cancer by miR-4319/SPNS2 axis (Jiang Y et al, 2021). HOXA-AS3 is significantly upregulated in oral squamous cell carcinoma(OSCC) and this high expression positively correlated with the pathological stage and poor prognosis of patients, which promotes the development of OSCC through sponging and inhibiting miR-218-5p (Zhao Y et al, 2021). Additionally, HOXA-AS3 knockdown reduced drug resistance by upregulating HOXA3 expression, and enable the development of novel and efficient strategies to treat non-small-cell lung carcinoma (Lin S et al, 2019). However, the role of HOXA-AS3 on androgen resistance in prostate cancer remains unclear. Therefore, we tried to find the potential targets of HOXA-AS3, aim to provide potential candidates for the early prevention and treatment of androgen resistance in prostate cancer in the future.

In this study, we revealed the expression pattern and role of lncRNA HOXA-AS3, and further explored its regulatory mechanism in promoting prostate cancer progression. The research mechanism is shown in Figure 1.

**Figure 1:**
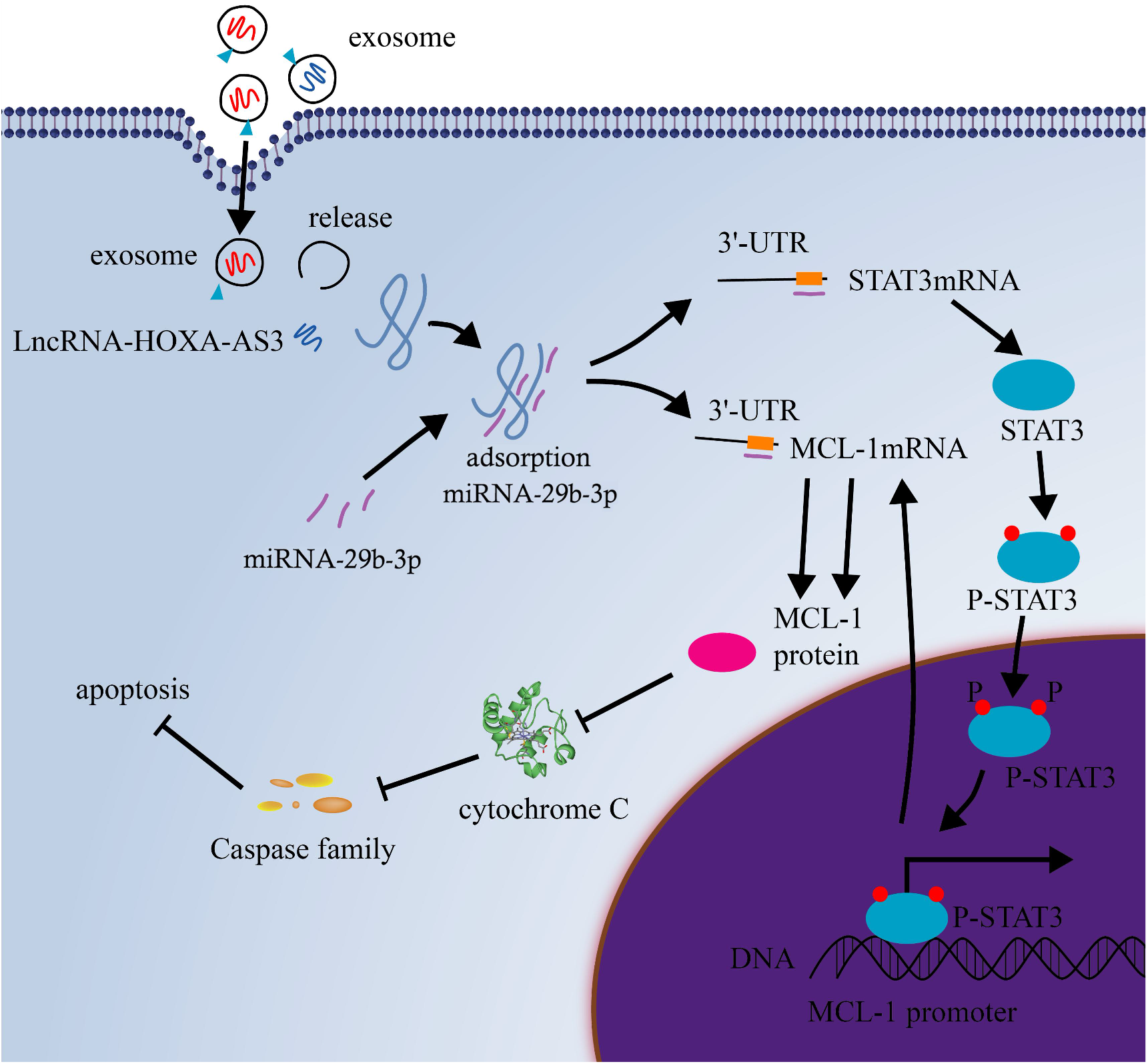
Schematic of the action of Exosome-derived LncRNA HOXA-AS3 regulating castration resistance and progression of prostate cancer.

## Results

### PC-3 cell-derived exosomes promote castration resistance in LNCaP cells

In order to clarify the transformation from ADPC to CRPC, ADPC cell model (LNCaP) were co-cultured with CRPC cell model (PC-3) for four weeks. We observed that the morphology of LNCaP cells changed significantly under light microscope, the cell mobility became stronger and migrated outward, the morphology was spindle shaped, became longer and the cytoplasm decreased significantly (Figure 2A). MTT results showed that in general medium, the proliferative ability of LNCaP cells co-cultured with PC-3 cells was not significantly changed compared with normal LNCaP cells; However, in androgen-deprived medium the proliferation of LNCAP cells was significantly increased after co-culturing (*P*<0.05)(Figure 2B). Clone formation ability of LNCaP cells co-cultured with PC-3 cells was stronger than that of normal LNCaP cells under both in general medium and androgen-deprived medium. In androgen-deprived medium, the difference between the two groups was more significantly (Figure 2C). We then separated the exosomes of PC-3 cell line by ultracentrifugation, and identified the extracted exosomes by transmission electron microscope. The transmission electron microscope results showed that the extracts had typical exosome characteristics(Figure 2D).We observed that exosomes derived from prostate cancer cell lines all expressed exosomes surface-specific markers TSG101, Alix, HSO70 in Western blot experiments(Figure 2E), and were successfully taken up by LNCAP and PC-3 cells (Figure 2F). Then we compared the proliferation of LNCaP cells with PC-3 exosomes and normal LNCaP cells by MTT. The results showed that the proliferative ability of LNCaP cells treated with PC-3 exosomes was significantly higher than that of normal LNCaP cells in general medium and androgen-deprived medium (P<0.05)(Figure 2G). The results of Colony formation assay showed that the clonogenic ability of LNCaP cells treated with PC-3 exosomes was stronger than that of normal LNCaP cells in general medium and androgen-deprived medium (P<0.05)(Figure 2H). Above all, we can get PC-3 cells promote androgen resistance of LNCaP cells through exosomes.

**Figure 2.**
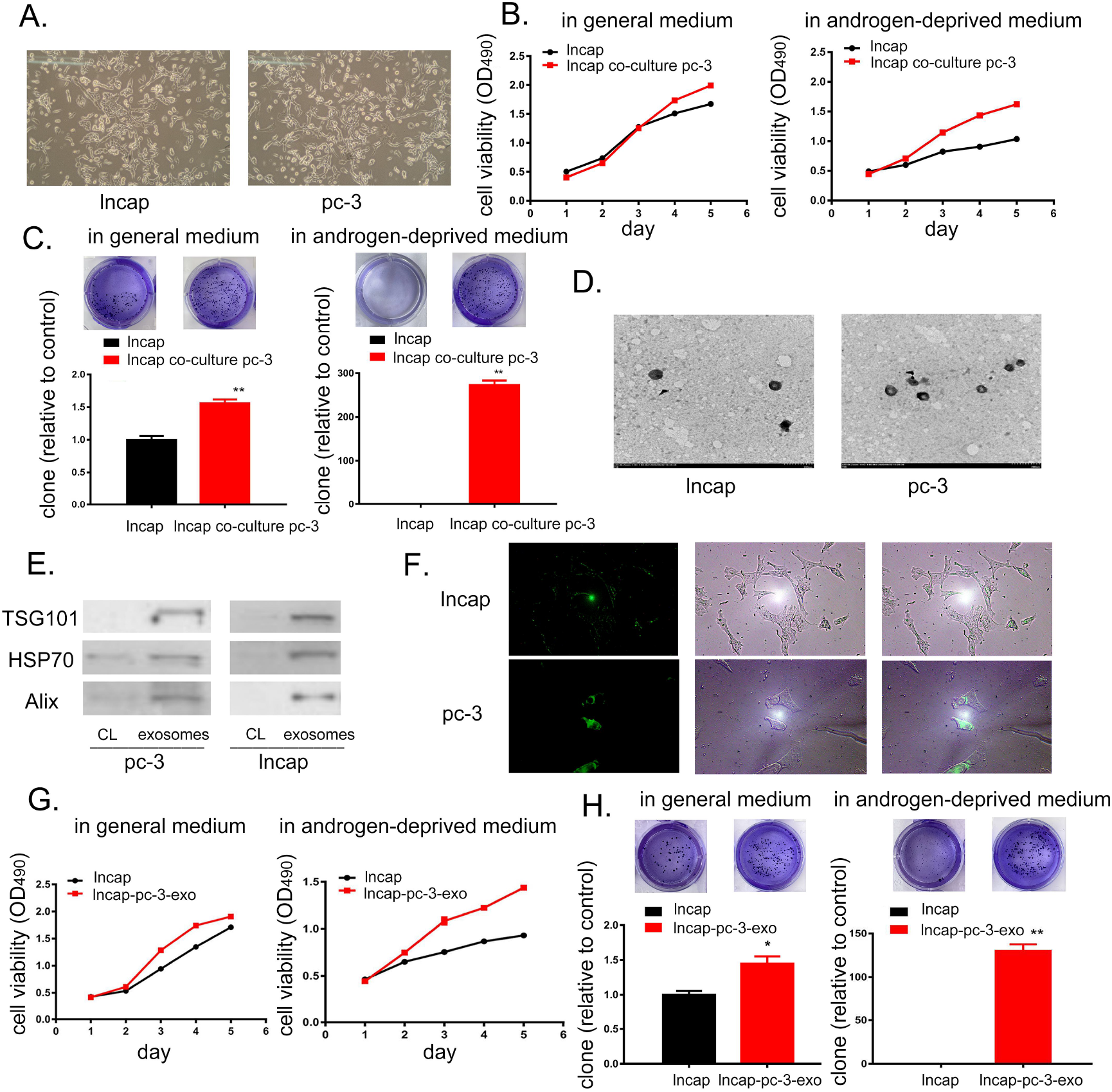
PC-3 cell-derived exosomes promote castration resistance in LNCaP cells. A. Morphology of LNCaP cells before and after co-culture (left: before co culture; right: after co culture)(200×); B. Cell activity of LNCaP cells cultured in general medium; Activity of co-cultured LNCaP cells in androgen-deprived medium; C. Clone formation ability of LNCaP cells co-cultured with PC-3 cells and LNCaP cells under general medium and androgen-deprived medium; D.Identification of exosome by transmission electron microscopy; E.Identification of exosome by WB; F. LNCAP cell and PC-3 cell uptake exosome(200×); G.Cell viability of LNCaP and LNCaP-PC-3-exo in general medium and androgen-deprived medium; H. The Monoclonal forming ability of LNCaP and LNCaP-PC-3-exo in general medium and androgen-deprived medium. \**P*<0.05, * \**P*< 0.01, \*\*\**P*< 0.001

### Exosomal HOXA-AS3 promotes castration resistance of prostate cancer cells

The differentially expressed lncRNAs in the exosomes of normal prostate (PNT) and cancer cells (LNCaP, cw22rv1, DU145, PC-3, VCAP) were obtained through bioinformatical analysis, 331 differentially expressed exosomal lncRNAs were found(Figure 3A). Furthermore, the differentially expressed exosomal lncRNAs in androgen-dependent (LNCaP, CWR22RV1) and androgen-independent (DU145, PC-3, VCaP) cancer cells were compared, there were 915 differentially expressed lncRNAs (Figure 3B) and four significantly different lncRNAs, HSPA8P19, AC10416.1, HOXA-AS3, RPL12P38 (Figure 3C&D). LNCaP cells were stimulated with different concentrations of exosomes (50 ng/ml, 100 ng/ml, 200 ng/ml and 500 ng/ml). The results showed that with the increase of exosome, the proliferation ability and expression level of HOXA-AS3 in LNCaP cells were increased (Figure 3E&F). Then, we selected PC-3 cells with relatively high expression of HOXA-AS3 for HOXA-AS3 knockdown experiments, and LNCaP with relatively low expression of HOXA-AS3 for HOXA-AS3 overexpression experiments. The results show that knockdown of HOXA-AS3 can significantly inhibit the proliferation of PC-3 (P<0.05), and overexpression of HOXA-AS3 can significantly enhance the proliferation of LNCaP (P<0.01) (Figure 3G). Flow assay of apoptosis showed that knockdown of HOXA-AS3 in PC-3 cells promoted PC-3 apoptosis, while overexpression of HOXA-AS3 in LNCaP cells inhibited LNCaP apoptosis (Figure 3H). These results revealed that HOXA-AS3 was associated with androgen resistance in prostate cancer.

**Figure 3.**
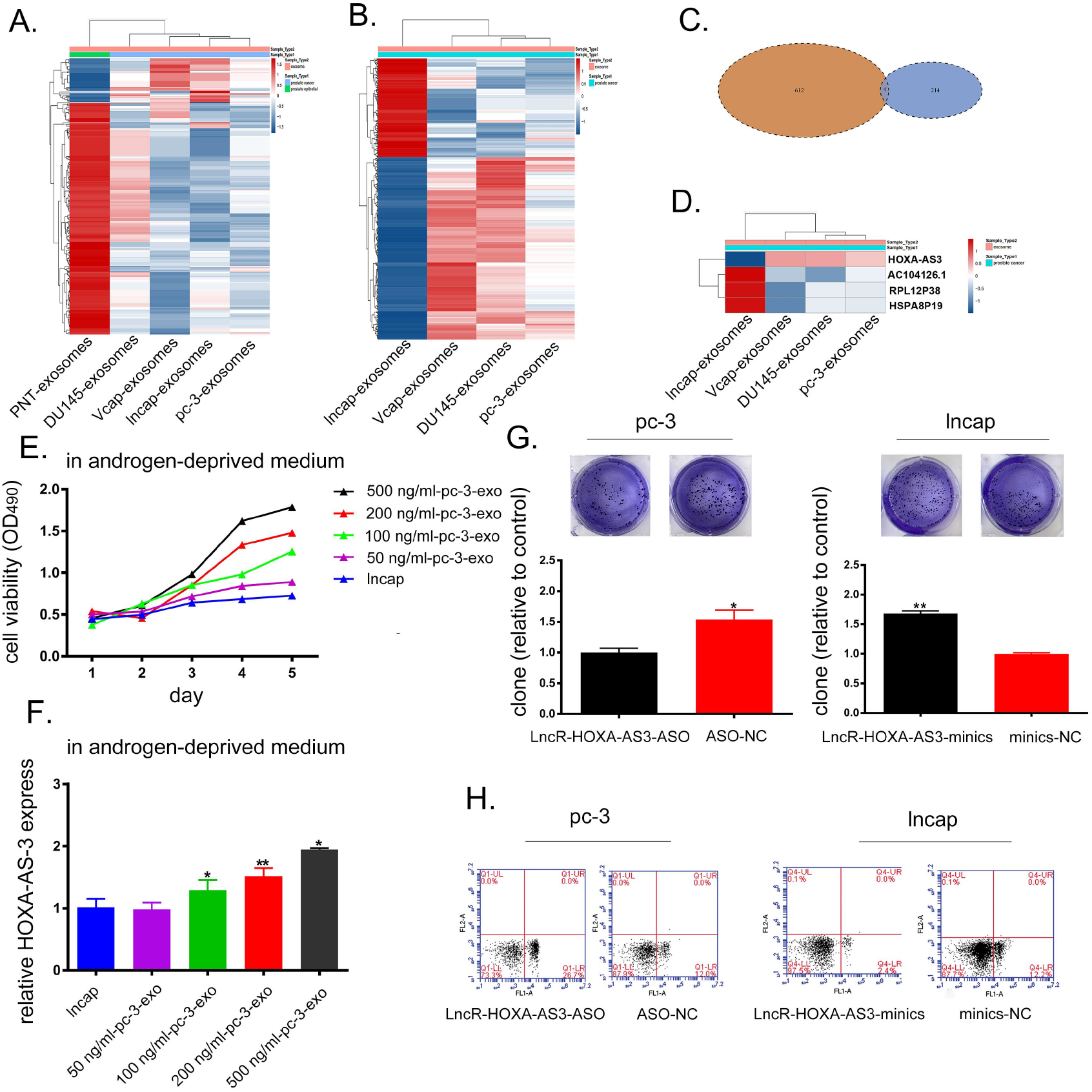
Exosomal HOXA-AS3 promotes castration resistance of prostate cancer cells. A. According to Shengxin analysis, lncRNAas differentially expressed in normal prostate (PNT) and cancer cell exosomes, red: high content; Blue: low content; B. Differentially expressed lncRNAs in exocrine of hormone-dependent (LNCaP, CW22RV1) and hormone-independent (DU145, PC-3, VCAP) cells of prostate cancer (red: high content; blue: low content); C. Wayne diagram of two types of lncRNA intersection; D. Intersection of differentially expressed lncRNAs (red: high content; blue: low content); E. Effects of different concentrations of PC-3 exosomes on the proliferation of LNCaP cells; F. Expression level of HOXA-AS3 content after LNCaP treated with different concentrations of exosomes; G. Knockdown of HOXA-AS3 reduced the colony-forming ability of PC-3, overexpression of HOXA-AS3 enhanced the colony-forming ability of LNCaP; H. Knockdown of HOXA-AS3 increased PC-3 apoptosis; overexpression of HOXA-AS3 decreased LNCaP apoptosis. \**P*<0.05, * \**P*< 0.01, \*\*\**P*< 0.001

### HOXA-AS3 as a competing endogenous RNA (ceRNA) regulates miR-29-3p to regulate tumor progression

The binding sites between HOXA-AS3 and miR-29b-3p were predicted using the lncBase and Starbase (http://starbase.sysu.edu.cn/). We then constructed HOXA-AS3-WT and HOXA-AS3-mut plasmids and co-transfected them with miR-29b-3p into HEK-293 cells respectively. Dual-Luciferase Reporter Gene Assays showed that HOXA-AS3-mut could not inhibit the luciferase level of miR-29b-3p, while HOXA-AS3-WT could inhibit the luciferase level of miR-29b-3p, suggested there was a targeting relationship between them (Figure 4A&B). In vitro experiments found that in PC-3 cells, overexpression of miR-29b-3p can inhibit the monoclonal formation and migration of PC-3 cells(Figure 4 C&E). At the same time, in LNCAP cells, knockdown of miR-29b-3p can promote LNCAP cell monoclonal formation and migration ability (Figure 4 D&F). These results revealed that HOXA-AS3 can specifically inhibit the expression of miR-29b-3p and affect the progression of prostate cancer.

**Figure 4.**
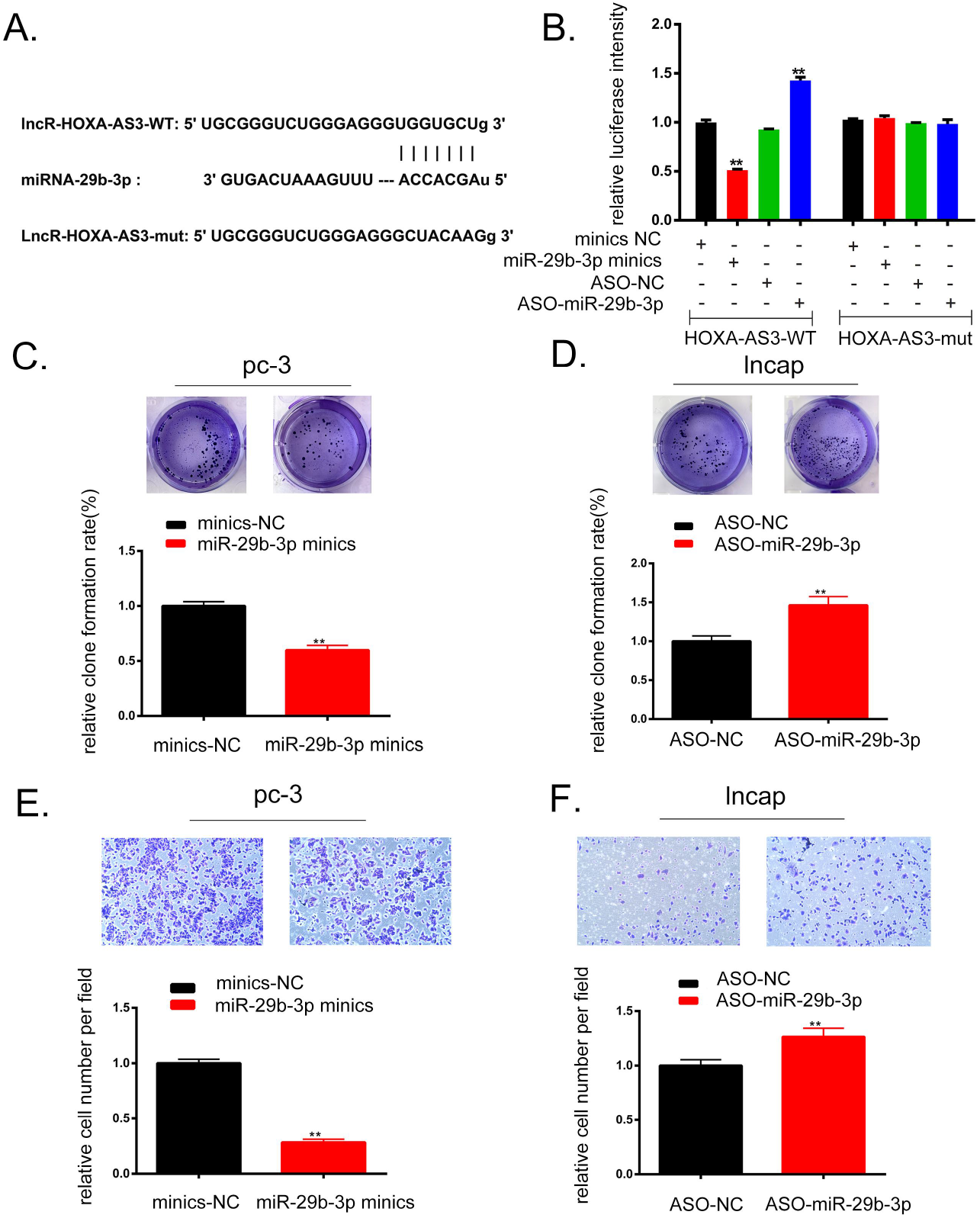
HOXA-AS3 as a competing endogenous RNA (ceRNA) regulates miR-29-3p to regulate tumor progression. A. Binding site mutation information; B. Dual-Luciferase Reporter Gene Assays to verify the targeting relationship between HOXA-AS3 and miR-29b-3p; C. Overexpression of miR-29b-3p reduces PC-3 cell colony formation ability; D. Knockdown of miR-29b-3p enhanced LNCaP cell colony-forming ability; E. Overexpression of miR-29b-3p reduces the invasive ability of PC-3 cells(200×); F. Knockdown of miR-29b-3p enhanced the invasive ability of LNCap cells(200×). \**P*<0.05, * \**P*< 0.01, \*\*\**P*< 0.001

### miR-29b-3p regulates the transcription and translation level of McL-1

Through the network database, we found miR-29b-3p can specifically bind to the 3’UTR region of STAT3/Mcl-1, and the specific binding site was shown in Figure 5A&B. Based on that, STAT3-WT, STAT3-mut, Mcl-1-WT, Mcl-1-mut plasmids are constructed respectively and co-transfected with miR-29b-3p into HEK-293 cells. STAT3-mut and Mcl-1-mut cannot inhibit the luciferase level of miR-29b-3p, while STAT3-WT and Mcl-1-wt can inhibit the luciferase level of miR-29b-3p, and both have a targeting relationship (Figure 5C&D). Then, Western blot and immunofluorescence experiments confirmed that overexpression of miR-29b-3p could inhibit Mcl-1, cytochrome c and caspases-9 levels (Figure 5E&F). On the contrary, knockdown of miR-29b-3p could promote Mcl-1, cytochrome c and caspases-9 levels (Figure 5G&H).

**Figure 5.**
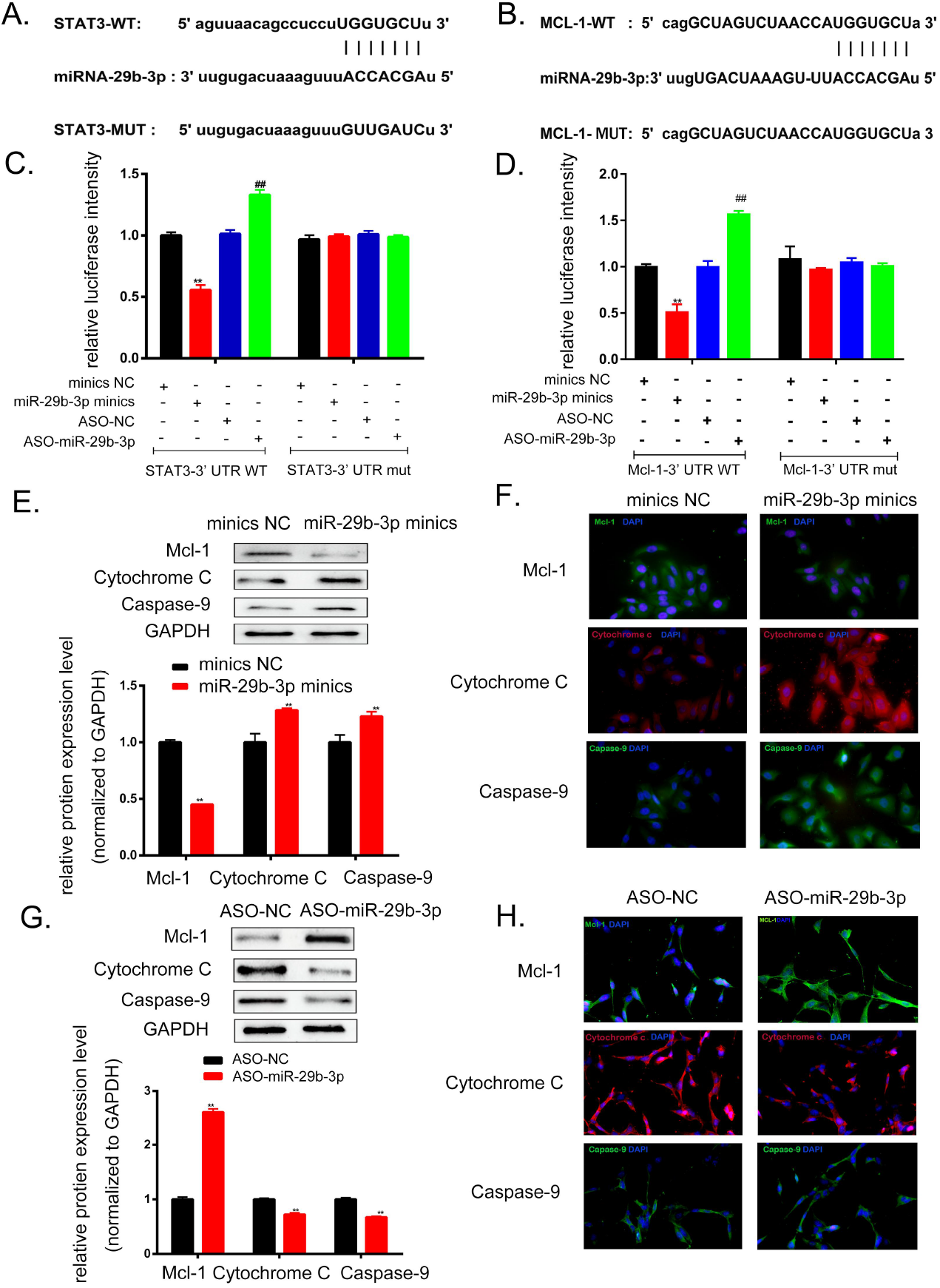
miR-29b-3p regulates the transcription and translatin level of McL-1. A. STAT3 and miR-29b-3p binding sites and mutations into site information; B. Mcl-1 and miR-29b-3p binding sites and mutations into site information; C. Dual-Luciferase Reporter Gene Assays to verify the targeting relationship between STAT3 and miR-29b-3p; D. Dual-Luciferase Reporter Gene Assays to verify the targeting relationship between Mcl-1 and miR-29b-3p; E. Expression of Mcl-1 and downstream proteins in PC-3 cells; F. Expression of Mcl-1 and downstream proteins in PC-3 cells immunofluorescence results(200×); G. Expression of Mcl-1 and downstream proteins in PC-3 cells; H. Expression of Mcl-1 and downstream proteins in PC-3 cells immunofluorescence results(200×); \**P*<0.05, * \**P*< 0.01, \*\*\**P*< 0.001

### STAT3 regulates the apoptosis, proliferation and migration of prostate cancer cells through Mcl-1/Cytochrome c/Caspase-9 pathway

The binding sites of STAT3 and Mcl-1 were predicted by JASPAR data, and Mcl-1-pomotor-WT and MCL-1-pomotor-mut plasmids were designed and transfected with STAT3 overexpression vector into LNCaP cells respectively to determine the relationship between STAT3 and Mcl-1. The transcriptional regulatory relationship of MCL-1. To eliminate the effect of miR-29b-3p on the experimental results, we knocked down the level of miR-29b-3p. Dual-Luciferase Reporter Gene Assays showed that STAT3 can activate the transcriptional level of Mcl-1 (Figure 6A&B). In colony experiments, overexpression of STAT3 in PC-3 and LNCAP can promote colony formation (Figure 6C). In cell migration assays, overexpression of STAT3 in PC-3 and LNCAP promotes the migratory capacity of cells (Figure 6D). Western blot experiment and immunofluorescence validation of STAT3 and downstream protein expression in PC-3 cells (Figure 6E&F). Western blot experiment and immunofluorescence validation of STAT3 and downstream protein expression in PC-3 cells (Figure 6G&H). Finally, it was confirmed that STAT3 could regulate the effect of signal axis on apoptosis, proliferation and migration of prostate cancer cells.

**Figure 6.**
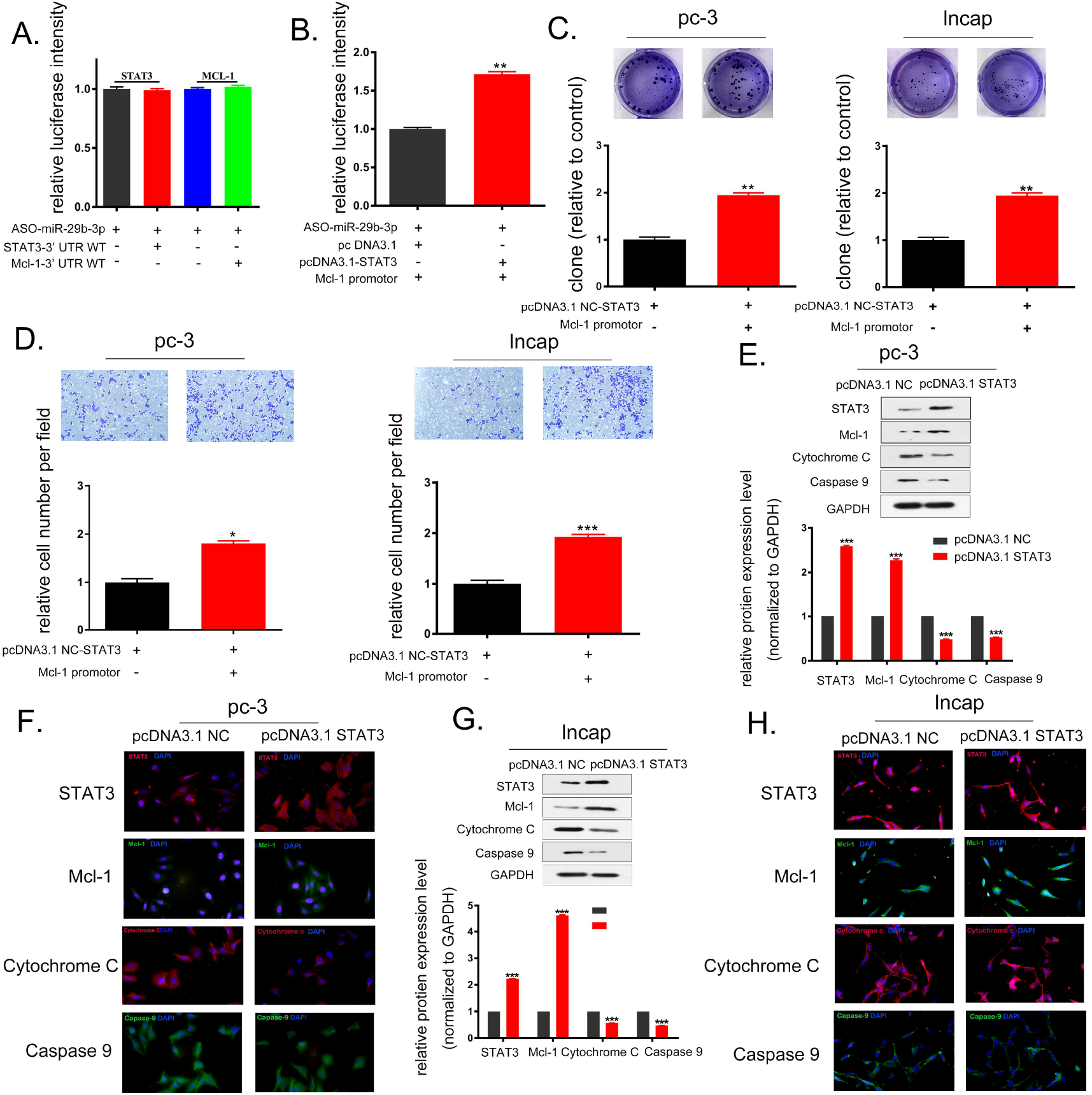
STAT3 regulates the apoptosis, proliferation and migration of prostate cancer cells through Mcl-1/Cytochrome c/Caspase-9 pathway. A. Dual-Luciferase Reporter Gene Assays verifies the fluorescein levels of STAT3 and Mcl-1 after interfering with miR-29b-3p; B. Dual-Luciferase Reporter Gene Assays verifies the binding of STAT3 and Mcl-1-promoter; C. Colony experiment verifies the proliferation ability of PC-3 and LNCaP cells; D.Transwell experiment verifies the migration ability of PC-3 and LNCaP cells(200×); E.Western blot verifies the expression of STAT3 and downstream proteins in PC-3 cells; F. Immunofluorescence detection of STAT3 and downstream protein expression in PC-3 cells(200×); G.Western blot verifies the expression of STAT3 and downstream proteins in LNCaP cells; H. Immunofluorescence detection of STAT3 and downstream protein expression in LNCaP cells(200×). \**P*<0.05, * \**P*< 0.01, \*\*\**P*< 0.001.

### HOXA-AS3 promotes castration resistance and progression of prostate cancer via miR-29b-3p/STAT3/McL-1/Cytochrome c/Caspase-9 axis

We transfected HOXA-AS3 and HOXA-AS3-ASO into LNCaP and PC-3 cells, respectively, and the corresponding untransfected PC-3 and LNCaP cells were served as controls. Exosomes were extracted and labeled as LNCaP-HOXA-AS3-exo, LNCaP-exo, PC-3-exo and PC-3-HOXA-AS3-ASO-exo, and identified by transmission electron microscopy (Figure 7A). The quantitative real-time PCR (qRT-PCR) experiments were performed to verify the expression levels of HOXA-AS3 in LNCaP-HOXA-AS3-exo, LNCaP-exo, PC-3-exo and PC-3-HOXA-AS3-ASO-exo exosomes, and the results showed that the HOXA-AS3 content in LNCaP-HOXA-AS3-exo was significantly higher than that in LNCaP-exo, while the HOXA-AS3 content in PC-3-HOXA-AS3-ASO-exo was significantly lower than that in PC-3-exo (Figure 7B). LNCaP cells were supplemented with LNCaP-HOXA-AS3-exo, LNCaP-exo and PC-3-exo, PC-3-HOXA-AS3-ASO-exo exosomes, and MTT was performed to verify the proliferation of LNCAP in androgen removal medium, the result showed that the proliferation of LNCaP cells after PC-3-exo exosomes stimulation was significantly higher than that of LNCaP cells stimulated by PC-3-HOXA-AS3-ASO-exo exosomes (P<0.05), while the proliferation of LNCaP cells after LNCaP-exo stimulation was significantly lower than that of LNCaP cells stimulated by LNCaP-HOXA-AS3-exo exosomes (P<0.05) (Figure 7C). Colony formation assay showed that the colony formation ability of LNCaP stimulated by adding PC-3-exo was significantly higher than that of LNCaP cells stimulated by PC-3-HOXA-AS3-ASO-exo exosomes, the colony-forming ability of LNCAP was significantly lower than that of LNCaP cells stimulated by LNCaP-HOXA-AS3-exo exosomes (Figure 7D). Western blot showed that the miR-29b-3p, cytochrome c and caspases-9 of LNCaP stimulated by PC-3-exo were lower than that of LNCaP cells stimulated by PC-3-HOXA-AS3-ASO-exo exosomes, while STAT3 and Mcl-1 were higher than LNCaP cells stimulated by PC-3-HOXA-AS3-ASO-exo exosomes (Figure 7E). In contrast, miR-29b-3p, cytochrome c and caspases-9 were higher in LNCAP stimulated by LNCaP-exo than LNCaP cells stimulated by LNCaP-HOXA-AS3-exo exosome, STAT3 and Mcl-1 were lower than LNCaP cells stimulated by LNCaP-HOXA-AS3-exo exosome(Figure 7G). We then verified the results by cellular immunofluorescence experiments and the results were consistent with Wb (Figure 7F&H). Thus, lncR-HOXA-AS3 promotes STAT3 and MCL-1 expression, inhibits cytochrome C release and Caspases-9 activation, and causes hormone resistance in LNCAP cells.

**Figure 7.**
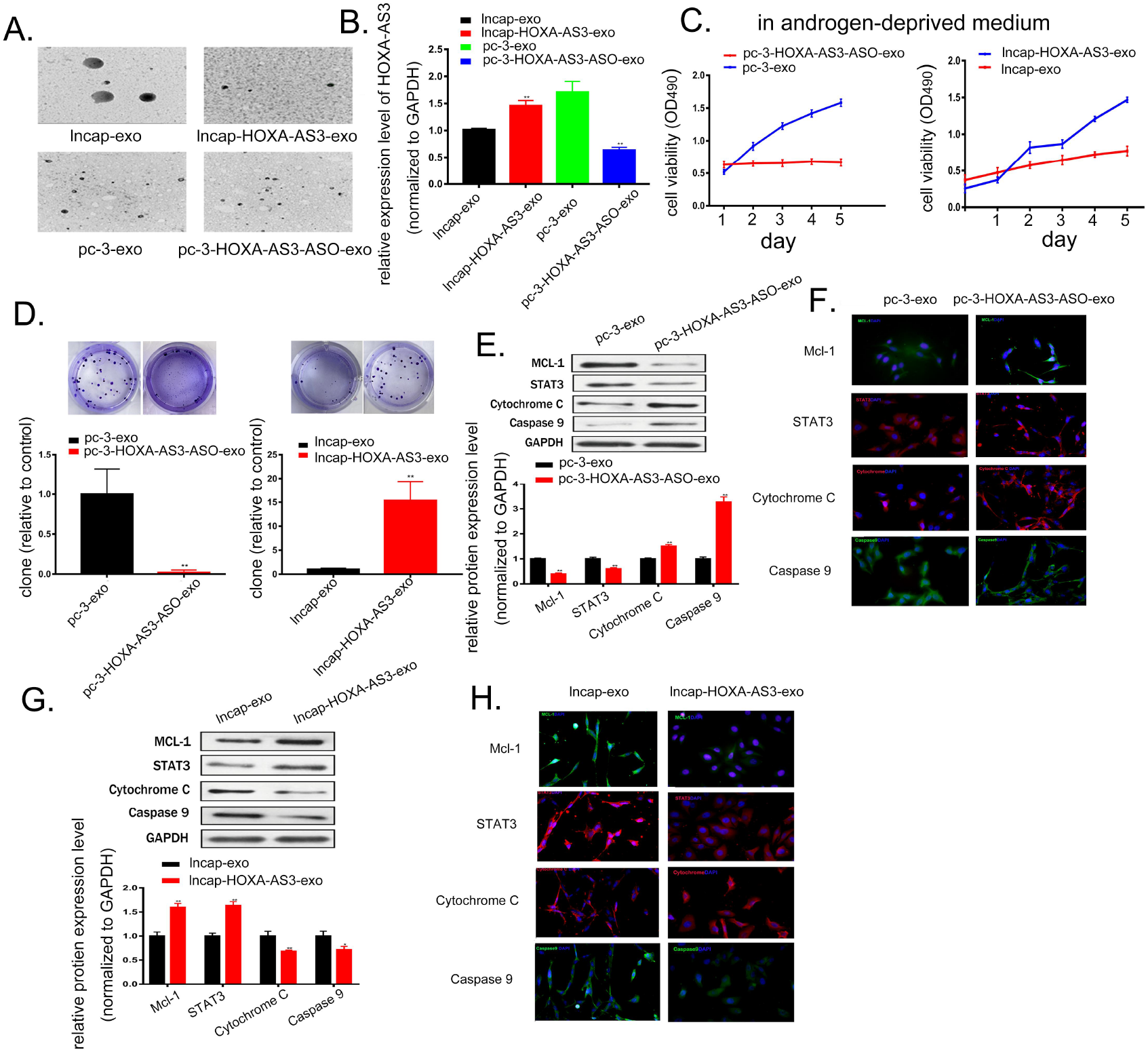
HOXA-AS3 promotes castration resistance and progression of prostate cancer via miR-29b-3p/ STAT3/McL-1/Cytochrome c/Caspase-9 axis. A. Identification of exosomes by transmission electron microscopy; B. Qpcr experiments identify the levels of HOXA-AS3 in exosomes in each group; C. MTT assay verify the proliferation of LNCaP after different exosome stimulation; D. Colony assay verify the formation ability of LNCaP cells after the addition of PC-3-exo and PC-3-HOXA-AS3-ASO-exo; Colony assay verify the formation ability of LNCAP cells after the addition of LNCaP-exo and LNCaP-HOXA-AS3-exo; E. Western blot assay of STAT3 and downstream protein gene expression in LNCaP after addition of PC-3-exo and PC-3-HOXA-AS3-ASO-exo; F. Immunofluorescence detection of STAT3 and downstream protein gene level expression after addition of PC-3-exo and PC-3-HOXA-AS3-ASO-exo(200×); G. Western blot assay of STAT3 and downstream protein gene expression in LNCaP after addition of LNCaP-exo and LNCaP-HOXA-AS3-exo; H. Immunofluorescence detection of STAT3 and downstream protein gene level expression after addition of LNCaP-exo and LNCaP-HOXA-AS3-exo(200×). \**P*<0.05, * \**P*<0.01, \*\*\**P*< 0.001.

### Exosomal HOXA-AS3 promotes hormone resistance and progression in prostate cancer was verified in vivo

We randomly divided the mice into two groups with five mice in each group. Then injected LNCAP cells into mice, and when the tumor reached 5-7 mm, the mouse testis was surgically removed, and PC-3-HOXA-AS3-ASO-exo exosomes and PC-3-exo exosomes were injected into mouse fluid every four days, tumor size was measured using vernier calipers every 3 days. The results showed that after androgen deprivation, the tumor size of mice treated with PC-3-exo increased slowly. On the contrary, the tumor of mice treated with PC-3-HOXA-AS3-ASO-exo was controlled, and the control rate reached 50% (Figure 8A&B&C). HE staining showed that apoptosis occurred in the tumor of mice treated with PC-3-HOXA-AS3-ASO-exo exosomes (Figure 8D). Wb, immunohistochemistry and immunofluorescence experiments confirmed that STAT3 and Mcl-1 in mice treated with PC-3-HOXA-AS3-ASO-exo exosomes decreased significantly, while caspases-9 and cytochrome c increased significantly, and the proportion of tumor cell apoptosis increased (Figure 8E&F&G).

**Figure 8.**
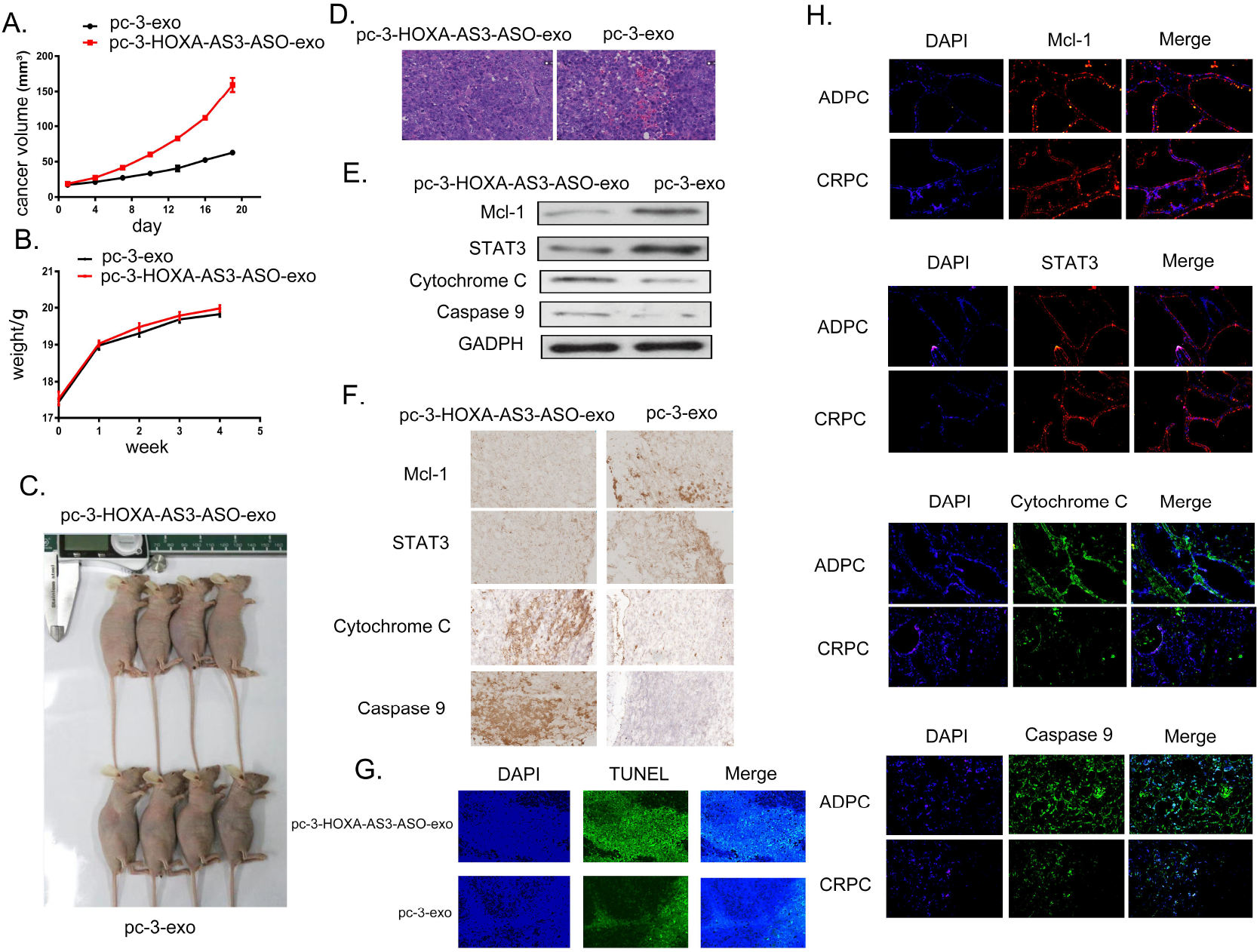
Exosomal HOXA-AS3 promotes hormone resistance and progression in prostate cancer was verified in vivo. A. Tumor volume in PC-3-exo and PC-3-HOXA-AS3-ASO-exo groups; B. Weight of nude mice in the PC-3-exo and PC-3-HOXA-AS3-ASO-exo groups; C. PC-3-exo and PC-3-HOXA-AS3-ASO-exo composition tumor situation; D. Pathology of HE in PC-3-exo and PC-3-HOXA-AS3-ASO-exo groups; E. Western blot assay of Mcl-1, STAT3, Cytochrome c, Caspase9 expression levels in PC-3-exo and PC-3-HOXA-AS3-ASO-exo groups(200×); F. Immunohistochemical detection of Mcl-1, STAT3, Cytochrome c, Caspase9 expression levels in PC-3-exo and PC-3-HOXA-AS3-ASO-exo groups(200×); G. TUNEL fluorescence staining of tumors in PC-3-exo and PC-3-HOXA-AS3-ASO-exo groups(200×); H. Immunofluorescence detection of MCL-1, STAT3, Cytochrome C, Caspase9 expression levels in ADPC and CRPC patients(200×).

Furthermore, we collected three cases of ADPC and three cases of CRPC in our hospital, and detected the expression of STAT3, Mcl-1, caspases-9 and cytochrome c in prostate cancer tissues by immunofluorescence. The results showed that the levels of STAT3 and Mcl-1 in ADPC were significantly lower than those in CRPC, and the levels of caspases-9 and cytochrome c were significantly higher than those in CRPC. These results were shown in Figure 7H which was consistent with the animal level.

## Discussion

During the transition from androgen-dependent to androgen-resistance prostate cancer, ADPC and CRPC cells may exist at the same time. We co-cultured LNCAP cells with PC-3 cells, it was found that the proliferation ability of LNCAP cells was enhanced, and the androgen dependence was reduced. Therefore, we are more convinced that the tumor microenvironment plays an important role in the transformation of prostate cancer from androgen-dependent to androgen-independent. Exosomes derived from cancer cells have a strong capacity to modify both local and distant microenvironments, play important functions in cancer development, metastasis, and anti-tumor or tumor-promoting immunity (Zhang L et al, 2019). Accumulating evidence suggests that exosomes play an essential role in the progression of prostate cancer. Yiming Zhang et al reported that androgen-independent prostate cancer (AIPC) cells promote the transition of ADPC cells into androgen-independent cells via exosome-mediated activation of HMOX1 (Zhang Y et al, 2021). Exosomes secreted by prostate cancer cells can promote the differ-entiation of bone marrow mesenchymal stem cells into myofibroblasts, which promote tumor proliferation and invasion (Chowdhury R et al, 2015). These results are consistent with the results of LNCAP/PC-3 cell co-culture experiments, and also confirmed that exosomes promote the transition of ADPC to CRPC.

lncRNAs have shown critical roles in biological processes, aberrantly expressed lncRNAs are strongly associated with the abnormalities in cell proliferation, migration, invasion, and apoptosis of multiple cancers (Lin W et al, 2020). By bioinformatic analysis, we found 4 lncRNAs that were significantly differentially expressed in ADPC and CRPC, among them, HOXA-AS3 is related to the occurrence and progression of various tumors. HOXA-AS3 facilitates the malignancy in colorectal cancer by miR-4319/SPNS2 axis. HOXA-AS3 was activated in lung adenocarcinoma and supported cancer cell progression (Zhang H et al, 2018). HOXA-AS3 could accelerate the growth of PC cells partially by regulating the miR-29c/CDK6 axis (Zhang X et al, 2021). However, the function of HOXA-AS3 in the development of androgen insensitive prostate cancer from androgen sensitive prostate cancer has not yet been determined. In the present study, the content of HOXA-AS3 in LNCAP cells increased with PC-3 exosome-stimulating concentration. Further, in PC-3 cells, knockdown of HOXA-AS3 significantly inhibited the proliferation of PC-3, while in LNCaP cells, overexpression of HOXA-AS3 significantly enhanced the proliferation of LNCaP. Subsequent endogenous validation and animal experiments also showed the same results, which confirmed the critical regulatory role of HOXA-AS3 in prostate cancer progression.

MiRNAs are endogenous small non-coding RNAs with the capacity to regulate gene expression post-transcriptionally. MiRNAs can conduct signal transduction with adjacent cells in the same or adjacent tissues through gap junctions to regulate cells (Rajagopal C et al, 2018). MiRNAs deletion will stimulate cell transformation and tumorigenesis in vivo (Kumar MS et al, 2007), which means that changes in miRNAs not only lead to tumorigenesis, but also promote the development of cancer. Recently, numerous studies have demonstrated that aberrant expression of miR-29b is common in the majority of human cancers. miR-29b is known to critically affect cancer progression by functioning as a tumor suppressor (Yan B et al, 2015). LncRNAs act as CeRNAs to regulate the biological function or expression of micRNAs (Zhang N et al, 2019; Yao N et al, 2019). Through bioinformatics analysis, we found that lncR-HOXA-AS3 can specifically adsorb microRNA-29b-3p, which was confirmed by the results of Dual-Luciferase Reporter Gene Assays. Further, in the subsequent overexpression and knockdown microRNA-29b-3p experiments, the results show the role of microRNA-29b-3p in the proliferation of prostate cancer cells was opposite to that of lncR-HOXA-AS3. This also confirmed that lncR-HOXA-AS3 could competitively adsorb micoRNA-29b-3p and thus influence the regulation of prostate cancer progression.

Myeloid leukemia-1 (Mcl-1) is an important anti-apoptotic protein mainly distributed in mitochondria and cytoplasm (Krajewski S et al, 1995), and its overexpression is associated with the pathogenesis and drug resistance of several cancers (Pan R et al, 2015; Gröbner SN et al, 2018). Given the anti-apoptotic properties of MCL-1, MCL-1 has been shown to be an intrinsic and acquired resistance factor that impairs the efficacy of various antitumor drugs (Wu W et al, 2020). As a regulator of MCL-1 expression (Jo MJ et al, 2019)., signal transducer and activator of transcription 3 (STAT3) is over-activated in tumor cells and is usually associated with poor clinical prognosis. STAT3 is involved in a variety of biological processes, including cell proliferation, survival, differentiation, and angiogenesis (Hanlon MM et al, 2019; Rawlings JS et al, 2004). STAT3 accelerates the proliferation and invasion of cancer cells and induces chemotherapy resistance (Hong H et al, 2021). In this study overexpression of micoRNA-29b-3p can significantly inhibit the level of STAT3/MCL-1 protein and the ability of monoclonal formation and migration of tumor cells, at the same time STAT3 activates the transcript level of MCL-1. The activation of MCL-1 by STAT3 inhibits the release of cytochrome C, which in turn inhibits the activation of Caspases-9, which leads to the decrease of cell hormone sensitivity and eventually leads to drug resistance.

In the immunofluorescence detection of prostate cancer tissue sections, STAT3 and Mcl-1 in ADPC patients were significantly lower than those in CRPC patients, and the levels of caspases-9 and cytochrome were significantly higher than those in CRPC patients. This result further confirmed the previous conclusion.

In view of the universality, expression specificity and functional importance of lncRNAs, lncRNAs have great potential as new biomarkers and therapeutic targets. At present, we have just begun to deeply study the biogenesis and functional mechanism of lncRNAs, and understand their clinical application value. With the continuous progress of sequencing technology and the development of new molecular and cell biology tools, the research field of prostate cancer lncRNAs will become a research hotspot. Undeniably, lncRNAs found in prostate cancer, combined with genome-wide functional screening, will produce a large number of clinically operable prostate cancer lncrnas.

In conclusion, exosomes play an important role in the transformation of prostate cancer from ADPC to CRPC. LncRNA HOXA-AS3 regulates the miRNA-29b-3p/Mcl-1/STAT3/cytochrome c/caspases-9 signaling pathway, thereby affecting the progression of prostate cancer. Therefore, lncRNA HOXA-AS3 can be considered as a promising intervention target to prevent the progression of prostate cancer, and also a potential target for the treatment of prostate cancer.

## Materials and methods

### Materials

The normal prostate (PNT) and cancer cell lines (LNCaP, CW22RV1, DU145, PC-3, VCaP) were supplied by the Cell Bank of the Chinese Academy of Sciences (Shanghai, China). miR-29b-3p-minics, miR-29b-3p-ASO, HOXA-AS3, HOXA-AS3-ASO, and its negative control small molecule RNA (miR-29b-3p-control, HOXA-AS3-control) were supplied by Guangzhou Provided by Ribo Biotechnology Co., Ltd. HOXA-AS3-WT, HOXA-AS3-mut, STAT3-WT, STAT3-mut, Mcl-1-WT, Mcl-1-mut plasmid, Mcl-1-pomotor-WT and MCL-1-pomotor-mut plasmid, pcDNA3. 1-STAT3 overexpression plasmids are all synthesized by BGI. PCR Master Mixkit was purchased from TaKaRa Company. Mcl-1 antibody, STAT3 antibody, Cytochrome c antibody, Caspase-9 antibody, HRP labeled fluorescent secondary antibody, TRITC and FITC labeled fluorescent secondary antibody were all purchased from Abcam (USA). TUNEL apoptosis kit and experimental animals were purchased from Beijing Weitong Lihua, and other cell culture consumables were purchased from Corning(USA).

### Cell culture

The prostate cancer cell lines LNCAP and PC-3 were supplied by the Cell Bank of the Chinese Academy of Sciences (Shanghai, China), and had been authenticated by Short Tandem Repeat (STR) profiling. These cells were maintained in RPMI-1640 supplemented with 10% fetal bovine serum (FBS, Gibco, USA) and penicillin (100 units/mL) and streptomycin (100 mg/mL) (Life Technologies) at 37°C with 5% CO_2_. Different media needed for the experiment were as follows: Normal medium (RMP1640+10% fetal bovine serum); Non-androgen medium (RMP1640+10% activated carbon/glucan treated fetal bovine serum); No exocrine medium (RMP1640+10% has no exocrine fetal bovine serum); Exosome-free androgen free medium (RMP1640+10% activated carbon/dexan treatment with excluded fetal bovine serum).

### RNA extraction and qPCR detection

After washing, cells and tissue samples were directly added with Trizol for RNA extraction. After RNA concentration and quality determination, take 2 μg RNA was reverse transcribed to obtain cDNA. According to the instructions of PrimeScriptTM RT Master Mix kit(manufacturer’s product number) and SYBR-Green PCR Master Mixkit (manufacturer’s product number), carry out RNA reverse transcription and qPCR. The relative mRNA expression was calculated using the 2-ΔΔCt normalized to GAPDH. The primers used in this study were as followed:

miR-29b-3p: Forward, GCCCAAAGGTGAATTTTTTGGG,
and reverse, CAGTGCGTGTCGTGGAGT.
U6: Forward, CTCGCTTCGGCAGCACA,
and reverse, AACGCTTCACGAATTTGCGT.
Mcl-1: Forward, CCCGTCCGTACTGGTGTTA,
and reverse, CGCGAGGCTGCTTTTCT.
STAT3: Forward, GGTTCAGCACCTTCACCMTT,
and reverse, CCCCGCACTTTAGATTCAT.
Cytochrome c: Forward, TGCTTGCCTCCCTTTTCA,
and reverse, ATGTCCCCCCGCACTTT.
Caspase-9: Forward, GGATGACCACCACAAAGCA,
and reverse, AATGGGACTCACAGCAAAGG.
GAPDH: Forward, TGACCCTTTTGGCTCCC,
and reverse, AAATCCCATCACCATCTTCC.

### MTT

MTT assays were performed using MTT Cell Proliferation Assay Kit (Cayman Chemical), following manufacturer’s instructions. After treatment, count and adjust the concentration of cells in each group, 1×10^4^/100μL inoculated into sterile 96-well plate. After the cells adhered to the wall, the cells at each time point were collected successively (24h, 48h, 72h, 96h and 120h). Before detection,add 20 μL of MTT reagent, and after incubation in the dark for 4 h, the plate was read by a microplate reader, and the OD490 data was achieved, repeat the biology three times.

### miRNA transfection

24 hours before transfection, the cells in each group were inoculated with 2×10^5^ cells per well density on a 6-well plate, and the transfection mixture was prepared according to Lipofectamine 2000 instructions, corresponding to each group of transfected cells, the final concentration of 50nM. After transfection for 12 hours, the cells were digested for subsequent experimental analysis.

### Plate cloning experiment

The cells in each group were digested with 0.25% trypsin and made into single cell suspension. The cells were counted and the cell suspension was diluted in gradient multiple. Each group of cells were inoculated with 100-300 cells respectively, and then inoculated into the 6-well plate, and gently swayed cross to make the cells evenly distributed, then cultured at 37 °C and 5% CO_2_ for four weeks. When visible clones begin to appear in the petri dish, the culture is terminated. Suck the cell culture medium and soak it twice with PBS. The cells were fixed for 15 min by 4% polymethase. After dyeing with an appropriate amount of crystal violet for 20 min, rinse with tap water and dry with air. Finally, invert the plate, directly calculate the number of clones, and repeat biology for three times.

### Transwell assay

The cells of each group were seeded in a Transwell chamber coated with Matrigel at a density of 2.5×10^4^ cells per well, and the lower layer was 750 μL of 5% FBS+RPMI1640 medium. After washing and methanol fixation, the cells were stained with crystal violet, and the cells that had not passed through the upper membrane were carefully removed with a cotton swab, and the cells passing through were observed under microscope. Each group was set with 3 duplicate wells, and 5 fields of vision were randomly selected from each well. The number of cells passing through the field was counted, and the average number was taken as the number of cells passing through this well, and the number of cells passing through 3 replicate wells was taken for statistical analysis.

### Isolation of Exosomes

Extract and isolate the exosomes of LNCaP and PC-3 cells: the prostate cancer cells were plated in 20 mL of RPMI-1640 containing 10% FBS, 100 units/mL penicillin, and 100 mg/mL streptomycin on 15cm dishes. When the cells reached approximately 80% confluence, they were washed 3 times with phosphate-buffered saline (1×PBS) and were then cultured in exosome-depleted RPMI-1640 medium. After 48h, the culture supernatant was collected and centrifuged at 3000×g for 15 min at 4°C. Then, an Ultracel centrifugal filter device with a 3 kD molecular weight cutoff (Millipore, USA) was used to concentrate the supernatant via centrifugation in a swing-out rotor and 4000×g at 4°C. The concentrated supernatant was further treated with ExoQuick-TC (System Biosciences, USA) for final exosome isolation. The exosome pellet was dissolved in 500 μL of 1×PBS. The extracted exosomes were identified by Western blot, transmission electron microscopy and particle size experiments.

### Excretion uptake test LNCAP+PC-3-exo PC-3+LNCAP-lncRNA HOXA-AS3-exo

Exosome uptake Kit (PKH67 membrane, Runji biology). Exosome treatment: dye is preheated at 37 °C and kept away from light. After pkh67labelingdye vortex centrifugation, take 5 μL add 50 μL reactionbuffer, add 50 μL exosomes were incubated at room temperature for 5 min and kept away from light. Purification steps: remove the plunger and add 200 μL sterilized PBS, 50xg centrifuged for 90s, discard the eluent, repeat the above steps, and put the labeled 100 μL exosome labeling preparation was applied to the top of the column, then 50xg was centrifuged for 90s, the eluent was discarded, the column was put into a 1.5ml centrifuge tube, and 200 was added to the top μL sterilized PBS, 50xg centrifuged for 90s, and the eluent was collected. Nuclear staining: add an appropriate amount of DAPI aqueous solution to PBS to prepare 5-15 μg/ml DAPI solution, add 1/10 of the volume of DAPI solution into the cell culture medium, culture the cells at 37 °C for 10-20 min, and wash the cells twice with PBS.

### Western blot, immunofluorescence and immunohistochemistry

Cells were lysed in RIPA buffer (Beyotime, China) containing protease inhibitors (Complete Mini Ethylene Diamine Tetraacetic Acid-Free Protease Inhibitor Cocktail, Roche). The protein lysates were centrifuged, and the supernatants were collected for protein quantification with a Bicinchoninic Acid assay kit (Pierce, USA). Protein lysates were resolved with 8% to 10% sodium dodecyl sulfate polyacrylamide gel electrophoresis (Beyotime, China), and were electrophoretically transferred to nitrocellulose filter membranes (Millipore). The membranes were then probed with antibodies(1:1000) against MCL-1 (abcam, ab32087), STAT3 (abcam, ab68153), GAPDH (abcam, ab8245), HSP70 (abcam, ab2787), TSG101 (abcam, ab125011), Alix (abcam, ab275377), Cytochrome C (abcam, ab133504) and Caspase-9 (abcam, ab32539).GAPDH (1:800; Cell Signaling Technology) and β-actin (1:1000; Cell Signaling Technology) were used as loading controls. After incubation with horseradish peroxidase (HRP)–conjugated goat anti-rabbit immunoglobulin G or HRP-conjugated goat anti-mouse immunoglobulin G secondary antibodies, the proteins were observed with laser confocal fiber lens. Immune complexes were detected by ECL Western blotting substrate (Thermo Fisher) and developed by bio rad (bio rad gel doc XR +).

### TUNEL apoptosis experiment

4% paraformaldehyde was fixed for 15min, 1% Triton X-100 was permeated for 5min, and TDT enzyme reaction solution was prepared according to TUNEL Kit (Nanjing Kaiji kga7061): The dosage of each sample is 45 μL Equilibration Buffer with 1.0 μL biotin-11-dUTP and 4.0 μL TdT Enzyme, ready to use, pay attention to avoid light. Each sample was dripped with 50 μL TDT enzyme reaction solution, put into a wet box to avoid light, and 60 min was protected from light at 37 °C. Mix 5 μL Streptavidin-TRITC reagent with 45 μL Labeling Buffer of each sample, and calculate the required total amount. Each sample was dripped with 50 μL Streptavidin-TRITC labeling solution, light avoidance reaction 30min at 37 °C, PBS washing 3 times, each time 5min.The nuclei were re-stained with DAPI (Hoechst) staining solution and 15min was protected from light at room temperature, PBS washed 3 times, each time 5min. Observed under fluorescence microscope, the excitation wavelength is 543 nm and the emission wavelength is 571 nm. Each experiment was repeated three times independently for statistical analysis.

### Dual-Luciferase Reporter Gene Assays

Starbase database (http://starbase.sysu.edu.cn/index.php) and TargetScan database (http://www.targetscan.org/vert_72/). UCSC (http://www.noncode.org/cgi-bin/hgTracks?hgsid=4056776) is used to predict the binding sites of miR-29b-3p with HOXA-AS3, STAT3 or Mcl-1 and the binding sites of STAT3 with Mcl-1-pomotor.The luciferase gene relationship was verified by luciferase assay. HOXA-AS3, STAT3, Mcl-1 and Mcl-1-pomotor fragments were amplified and cloned into pMIR-REPORTM luciferase vector (Promega) to obtain HOXA-AS3-WT, STAT3-WT, Mcl-1-WT and Mcl-1-pomotor reporter vectors. HOXA-AS3, STAT3 or Mcl-1 fragments containing mutant binding sites were cloned into pMIR-REPORTM fluorescent enzyme vector and named as HOXA-AS3-mut, STAT3-mut or Mcl-1-mut reporter vector. Co-transfected these notification vectors with miR-29b-3p minics or NC into PCa cells by Lipofectamine-3000, and then incubated for 48 hours. Dual-Luciferase ^®^1000 Assay kit (Promega) is used to detect firefly luciferase activity.

### Animal experiment

This experiment was approved by the animal experiment Committee. Ten 6-week-old male BALB/c nude mice (weighing 18-22g) were from the experimental animal resource center of the Chinese Academy of Sciences (Beijing, China). The mice were randomly divided into two groups with 5 mice in each group. LNCaP cells were collected, suspended in PBS at a density of 1×10^7^ cells / ml, and injected subcutaneously into the right side of the back rib of mice. After the tumor diameter reached 5-7mm, testicles of male mice were surgically removed, PC-3-exo-sh-HOXA-AS3 or PC-3-exo-control complex was injected into the tumor every 4 days (28 days in total), and tumor size was measured with vernier caliper every 3 days. calculated volume by V = 0.5 L ×W2(W: short diameter; L: long diameter). On day 28, mice were all sacrificed for further analyses.

### Statistical analysis

Each experiment in this study was repeated three times. The data were calculated using GraphPadPrism7 software and expressed as mean ± SD. The differences between the two groups were assessed by Student’s t test. One way ANOVA and Bonferroni post hoc comparison test were used to analyze the differences between multiple groups. *P* values <0.05 were considered to be significant differences.

## Acknowledgements

We would like to thank Professor Yuwan Liu from Haihe Laboratory of Synthetic Biology for his valuable comments on the manuscript. We wished to sincerely thank HaiJie Hu of the Institute of Key Laboratory of Industrial Fermentation Microbiology, Ministry of Education, Tianjin Industrial Microbiology Key Laboratory, College of Biotechnology, Tianjin University of Science and Technology for critical technical support.

## Competing interests

The authors declare that they have no competing interests.

## List of abbreviations

PCa: Prostate cancer
ADPC: Androgen-dependent prostate cancer
CRPC: Castration resistant prostate cancer
lncRNAs: Long noncoding RNAs
ADT: Androgen-deprivation therapy
Mcl-1: Myeloid cell leukemia-1
STAT3: Signal transducer and activator of transcription 3
FBS: Fetal bovine serum
ceRNA: competing endogenous RNA.

